# Conservation of expression regulation throughout the animal kingdom

**DOI:** 10.1101/007252

**Authors:** Michael Kuhn, Andreas Beyer

## Abstract

Following the increase in available sequenced genomes, tissue-specific transcriptomes are being determined for a rapidly growing number of highly diverse species. Traditionally, only the transcriptomes of related species with equivalent tissues have been compared. Such an analysis is much more challenging over larger evolutionary distances when complementary tissues cannot readily be defined. Here, we present a method for the cross-species mapping of tissue-specific and developmental gene expression patterns across a wide range of animals, including many non-model species. Our approach maps gene expression patterns between species without requiring the definition of homologous tissues. With the help of this mapping, gene expression patterns can be compared even across distantly related species. In our survey of 36 datasets across 27 species, we detected conserved expression programs on all taxonomic levels, both within animals and between the animals and their closest unicellular relatives, the choanoflagellates. We found that the rate of change in tissue expression patterns is a property of gene families. Our findings open new avenues of study for the comparison and transfer of knowledge between different species.

## Introduction

Gene functions have traditionally been determined using molecular and cellular approaches involving forward or reverse genetics. Functional annotations that were directly determined through these approaches are, however, not available at all for most species, and incomplete even for model species [1]. For non-model species, often only data transferred from other organisms is available. In this case, the degree of conservation of functions is uncertain, especially when a gene is duplicated in a non-model species, but not in the model species where its function has originally been studied. Previously, gene coexpression data has been used to find conserved coexpressed modules [2, 3] and to uncover functional similarities between genes from different species [4]. However, the latter approach requires that the two species are well-studied in both gene expression and functional annotation, and will suffer from incomplete and biased annotations [1].

Tissue expression data is available for many species, as tissues can be gathered even from non-model species where genetic tools such as transgenesis or RNAi are not available. Developmental gene expression profiles between closely related species can be compared to find functional links between genes and to detect differences between orthologs [5, 6, 7]. For closely related species, homologous tissues can easily be identified [8], and cross-species correlations between equivalent tissues of closely related species have previously been investigated [9, 10, 11]. Existing approaches require that expression datasets have been obtained under comparable conditions for the respective species. Across larger evolutionary distances, only few clearly homologous tissues can be determined. Even between closely related species, the relative amounts of cell types within tissues may change. This reliance on homologous tissue is, therefore, a severe limitation for functional mapping between many species: it is not possible to correlate gene expression patterns across species using the traditional methods. If it was possible to compare expression patterns across large phylogenetic distances, we could substantially improve the annotation of non-model-species genomes, fill annotation gaps in model species, and in particular address the problem of functional conservation after gene duplications.

To ameliorate this situation, we have developed a method to map tissue expression patterns of genes from one species to another, without defining equivalent tissues between the two species. Our hypothesis is that groups of functionally related genes will be coexpressed in very different tissues and species due to the re-use of ancestral functional modules. For example, it is possible to identify deep homologies among tissues [12], like homologous structures in the nervous systems of vertebrates and annelids [13,14]. Other organs show functional convergence, e.g. mammalian liver and brown fat in flies, which both carry out xenobiotic clearance functions [15]. For each gene of the source species, our approach predicted a virtual tissue expression pattern in the destination species. The correlation between these virtual expression patterns and the actually observed expression could then be used to score how well a gene's expression of a gene in a target species can be predicted from the expression patterns of its orthologs. Importantly, this scheme can be used to determine the extent to which the transcriptional regulation of sets of genes is conserved across large phylogenetic distances.Subsequently we illustrate the potential of our modeling approach with two applications: determining the degree of conservation of tissue-specific gene expression patterns, and for comparing the speed of functional divergence between independently evolving members of protein families.

## Results

To analyze tissue expression across the entire metazoan kingdom, we gathered genome and tissue expression data from 36 datasets covering 27 different species (Table 1, Table S1). The datasets contained both developmental time courses (e.g. embryonic stages) and static measurements of different tissues (like adult organs; see Supplementary File 1 for a complete list). For the sake of brevity, we refer to the all of these samples as “tissues.” Datasets were imported and normalized per gene (see Methods), i.e. we quantified only relative expression changes of the same gene between tissues, instead of comparing expression differences of genes within the same tissue. (Therefore, housekeeping genes and other genes which are globally expressed did not skew ouranalysis.) When we applied the concept of looking for correlations between orthologs across species to an existing dataset [10], we found that many of the reported lineage-specific expression shifts only changed the absolute expression levels, while the relative expression patterns remained conserved (Fig. S1). Normalizing each gene’s expression individually also avoided technical concerns regarding the comparability of absolute expression values between genes. However, this gene-wise normalization means that the normalized values are influenced by the complement of tissues that have been measured. For this reason, we only include datasets that survey a whole organism or a wide range of developmental time points. The datasets excluded during quality control (see next section) have between five and ten data points. Therefore, six diverse tissues seemed to be a lower limit for the number of data points.

**Table 1:**
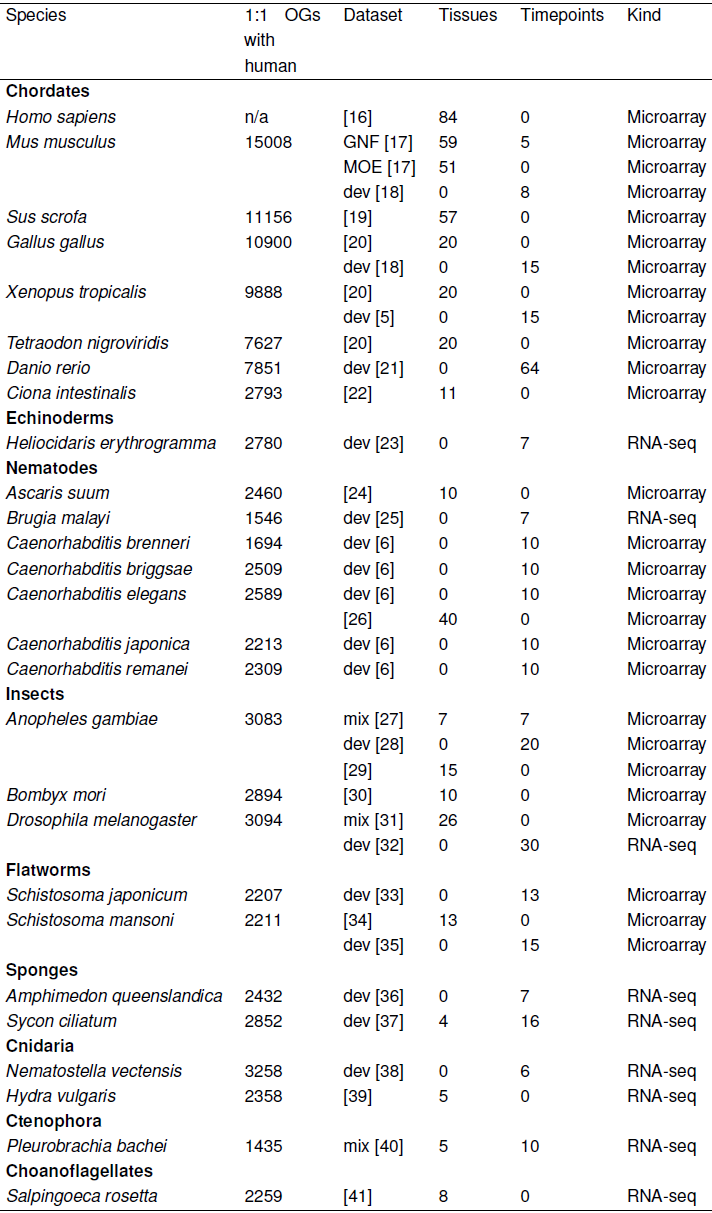
Analyzed species and datasets.

## Quality Control

The available datasets differ in their suitability for cross-species mapping. Existing measures for the quality of expression datasets rely on conserved features, e.g. conserved coexpression [42]. Because of the large biological diversity of species in our dataset, relying on conservation of features was not appropriate. We therefore devised a simple measure of dataset quality that only relied on the features of the given dataset. For each normalized dataset, we performed Principal Component Analysis and determined the proportion of variance represented by each eigenvector. We then calculated the fraction *v*_50_ of components that represent at least half of the total variance. For example, for the *C. elegans* dataset, the first four out of forty principal components explain just above 50% of the variance (hence, *v*_50_ = 0.1). Based on the observed correlation between *v*_50_ and median mapping quality (Fig. S2), we chose v_50_ ≥ 0.25 as a filter to remove the five worst datasets from our analysis: *Hydra vulgaris, Amphimedon queenslandica, Bombyx mori, Brugia malayi* and *Ascaris suum.*

## Mapping Gene Expression Between Species

In order to compare expression patterns across distant species, we first need to map the patterns. Our concept rests on the notion that the expression of a gene in a specific tissue of a target species can be predicted using the expression pattern of that gene across the tissues in the source species. For example, a gene specifically expressed in insect neurons is likely to be expressed also in the mouse brain. Here we show that this concept holds even if these “matching” tissues are not known. Consider the example of mapping gene expression patterns from fly to mouse (Fig. 1). We model *m_g_*,*_t_*, the relative expression of gene *g* in mouse tissue *t*, as a linear combination of the relative expression levels in all fly tissues (*f*,_*ĝs*_):

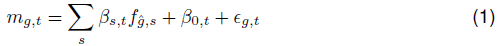

where *ĝ* is the fly ortholog of gene *g* and *ϵ* is the residual error. The regression coefficients *β_s_*,*_t_* and the intercept *β*_0_,*_t_* are fitted using all 1:1 orthologs between mouse and fly. Subsequently, this model can be applied to all fly genes to predict the expression in mouse tissue *t*. We used linear models in this first description of the method as they are a simple, transparent and efficient method that is relatively robust to over-fitting. Of course, other methods may be used as well. For example, Random Forest regression [43] can deal with non-linearity, while the lasso [44] could be used to deal with redundancy between source tissues.

**Figure 1:**
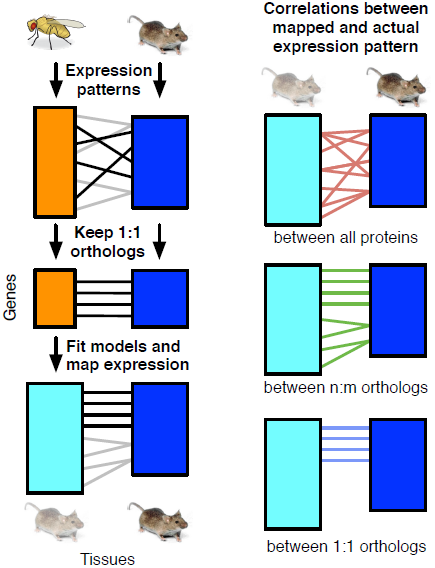
Mapping expression patterns across species. For each tissue in the target species, models were fitted to predict the tissue-specific gene expression pattern from the expression patterns of 1:1 orthologs in a source species. Mapping the expression patterns of all genes created virtual expression patterns, which could then be used to compute correlations between the mapped and actual expression patterns.

## Expression Distances Between Genes

After mapping expression patterns between species, we quantified how well a gene can be predicted by correlating its predicted expression across all tissues with the respective measurements of the target species using Pearson’s correlation coefficient (see Methods). These pairwise correlations between genes could be calculated for different sets of genes: phylogenetically unrelated genes, orthologs and 1:1 orthologs. Of these, 1:1 orthologs had the highest correlations (Fig. 2A). However, the overall distribution of correlations differed between dataset pairs, e.g. due to the varying number of tissues or variable data quality. Therefore, we computed an expression distance based on the quantiles *q_x_*,*_y_* of the matrix of correlation coefficients **R** = [*r_x_,_y_*]. We found that lineage-specific genes (i.e. those without homologs between the two species under consideration) tended to have lower correlations than genes with homologs. Therefore, to calculate quantiles, we computed the matrix R only for genes with homologs between the two species. We first analyzed correlations between 1:1 orthologs and checked if they depended on different properties of the genes (Fig. S3). We found that when the target gene had many coexpressed genes in the same species, the cross-species expression correlation tended to be higher (Fig. S3). To correct for this effect, we considered only target genes with similar numbers of coexpressed genes when computing the expression distance (Fig. 2 and Fig. 9 in Methods section). By design, the expression distance of background gene pairs had an uniform distribution. When making inferences about 1:1 orthologs, we used linear models based on a 10-fold cross-validation.

**Figure 2:**
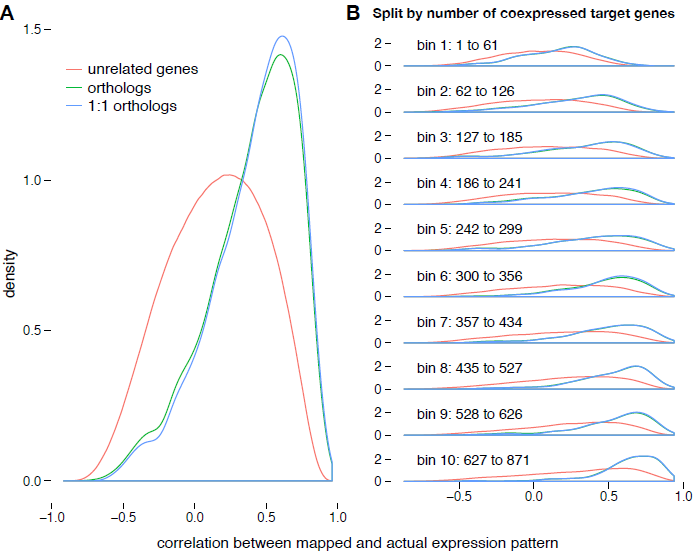
Distribution of correlations between mapped and actual expression patterns. When mapping expression patterns from fly to *C. elegans,* correlations between orthologs (green) and 1:1 orthologs (blue) were much higher than for background gene pairs (pairs of genes that are not homologous to each other, shown in red). (**b**) Target genes were split in bins according to the number of genes with similar expression patterns within the target species. Pairs of background genes had a higher correlation when there were more genes with similar expression patterns, as is evident from the shift towards higher correlations. For this pair of datasets, bins contained between 321 and 326 one-to-one orthologs, with an average of 324.

## Benchmarks

In order to establish the biological relevance of our expression distance measure, we applied benchmarks at three levels, namely sequence, structure, and function. On the sequence level, we found that expression distances could be used as a signal to decide which of the top two BLAST hits for a query protein is the true 1:1 ortholog of the query protein in the target species (Fig. 3A and Fig. S4). On the structural level [45], expression distance and the number of proteins belonging to a structural fold were correlated (Fig. 3B and Fig. S5). That is, structural folds with fewer members, and hence lower functional diversity, were more similar in their expression patterns across species.

**Figure 3:**
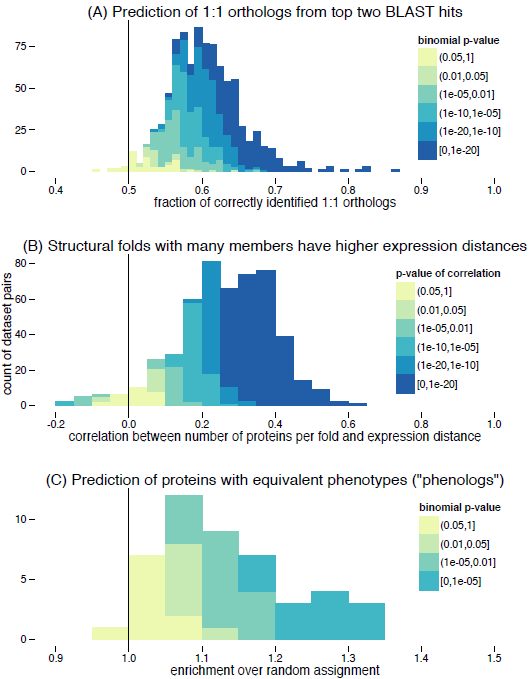
Summary of benchmarking results. For each pair of datasets from different species, the performance in different benchmarks has been computed, along with a p-value. In all three benchmarks, there was a clear shift of the results relative to the random expectation (black line). For details, see Fig. S4, Fig. S5, Fig. S6 and supplementary text. Due to limited structural and functional annotations, there was a lower number of dataset pairs for the two lower panels.

Lastly, on the functional level, we applied the phenolog concept [46] to find equivalent phenotypic annotations across species. We found that expression distances could be used to better predict which member of a protein family has been annotated with a matching phenotype (Fig. 3C and Fig. S6).

## Conservation of Gene Expression Programs

At all taxonomic levels, we determined the conservation of the expression patterns of 1:1 orthologs. This data then allowed us to estimate the degree of conservation of tissue-specific expression patterns, even between groups of species that do not have readily identifiable homologous organs. For each pair of datasets, we first computed the median expression distance of 1:1 orthologs (Fig. 4). We then tested for each pair of datasets whether the distribution of expression distances between 1:1 orthologs was shifted towards lower values, i.e. if the median is below 0.5. Using the Wilcoxon signed-rank test and controlling for multiple hypothesis testing with the Benjamini-Hochberg method [47], we found that all dataset pairs had significant shifts to lower expression distances (q < 0.05 and median < 0.5). This analysis revealed both an expected enrichment for closely related species and unexpectedly high enrichments between very distant species, such as between chordates and insects. In general, developmental datasets mapped less well to other species than datasets of adult tissues.

**Figure 4:**
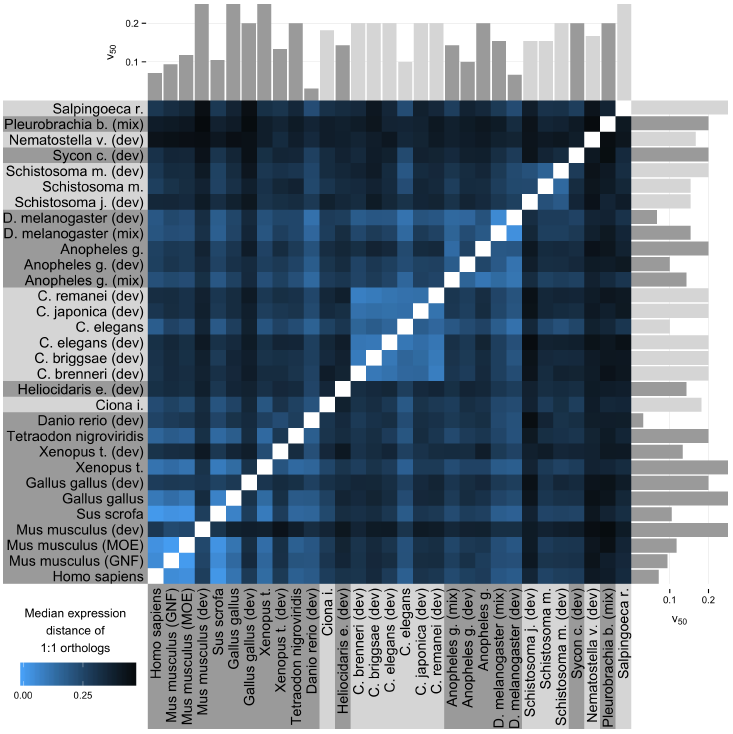
Conservation of expression patterns throughout the metazoans and choanoflagellates. For all dataset pairs, the median expression distance of 1:1 orthologs is shown. Within most clades, this median was very low and approached 0 in some cases. When there was no enrichment of 1:1 orthologs towards lower expression distances, the median was 0.5 (see Fig. 5). See Fig. S7 for a version of this figure without filtering for dataset quality.

To summarize the data shown in Fig. 4, we computed median expression distances for 1:1 orthologs across all internal nodes of the phylogenetic tree (e.g. for vertebrates, we compared expression patterns between fish and tetrapods). As the median expression distances vary greatly between dataset pairs, we also computed the distribution of expression distances and the number of well-conserved OGs for the best dataset pair across each internal node (Fig. S8 and Fig. S9). Using a Wilcoxon signed rank test, we then tested if the distribution of median expression distances is shifted towards lower values, i.e. if the median of the distribution is lower than 0.5. This was the case for all internal nodes, with the highest p-value (5e-49, median value: 0.39) observed when mapping from the ctenophore *Pleurobrachia* to other animals. This confirmed that our approach could predict expression patterns over large evolutionary distances (Fig. 5 and 6). For some clades, the available data was very uneven on the two sides of the internal node. For example, at the level of eumetazoa, only one species with few tissues was available for cnidarians, whereas most bilaterian species had many tissues measured. Thus, expression distances were higher when mapping from cnidarians to bilaterians than the other way round. Interestingly, the median divergence between animals and the outgroup choanoflagellates was comparable to the median divergence between major animal clades, e.g. bilateria. Thus, mapping tissue-specific gene expression revealed expression programs conserved for 1 billion years.

**Figure 5:**
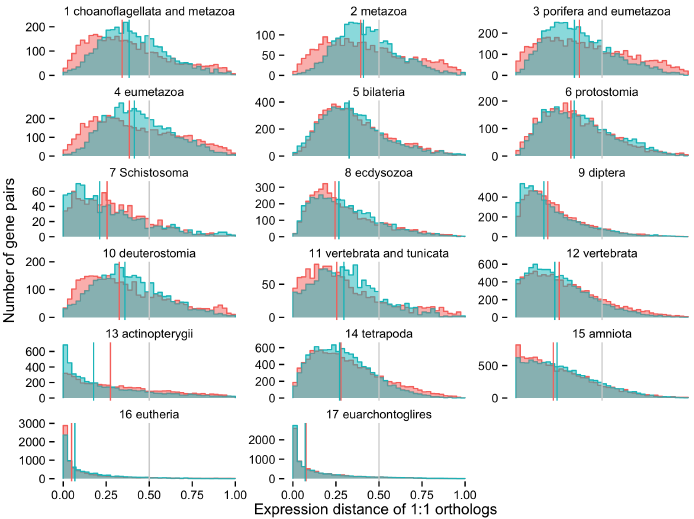
Distribution of median conserved expression. For each clade, the distribution of expression distances of 1:1 orthologs is shown. Red and blue colors denote the direction of the mapping, either from the first subclade to the second or vice versa. For each distribution, the median is shown as a vertical bar. The gray bar corresponds to an expression distance of 0.5, which is the median to be expected by chance. When the mapping is successful, our mapping procedure yields virtual expression patterns of 1:1 orthologs that are very similar to the actual expression patterns, and the distribution of expression distances is skewed towards lower values. Our mapping procedure becomes less accurate over larger evolutionary distances, and the distribution of expression distances becomes less skewed. It becomes a uniform distribution when 1:1 orthologs cannot be mapped better than background gene pairs. Clades are numbered corresponding to the taxonomic tree in Fig. 6.

## Correlations Between Expression Changes of Homologs

Next, we addressed the question if conservation of expression programs depends on the functions of genes, i.e. if certain gene functions generally imply a stronger conservation of expression programs than other functions. To this end, we compared the expression distances of gene families in different clades under the assumption that functional constraints would lead to expression conservation in independent clades. If the rate of expression divergence is a property of the gene family, we expect a correlation between the expression similarities for each family in different clades. In other words, a gene family that has a conserved expression pattern in one clade should also have a conserved expression pattern in another clade. For each internal node with two or more species on either side of the split, we calculated the median expression distance per gene family within each of the two clades. Out of four internal nodes with more than one species on both sides, we found significant Spearman correlations (r_s_) of median expression similarities for three splits (Fig. 7A): between tetrapods and fishes (r_s_=0.18, #12 in Fig. 5), between protostomes and deuterostomes (r_s_=0.15, #4), and between nematodes and insects (r_s_=0.06, #7).

**Figure 6:**
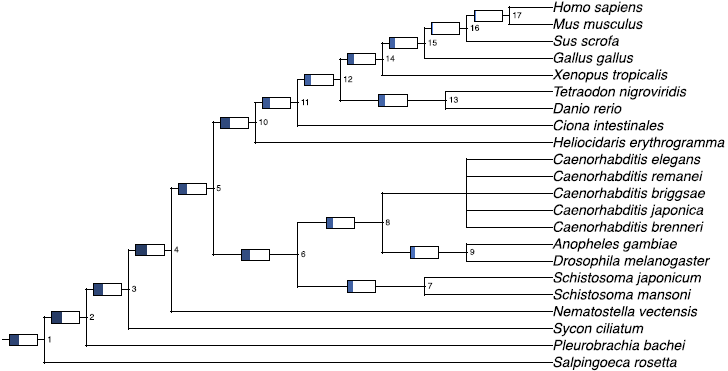
Conservation of expression patterns across clades. The length of the blue bars denotes the median expression distance of 1:1 orthologs across the bifurcation, with values between 0 and 0.5. (0 corresponds to the best possible value, while 0.5 would occur when there is no enrichment of lower expression distances.) The numbers next to the internal nodes refer to the clade numbers in Fig. 5

The previous analysis was only possible for a subset of the taxonomic splits in our body of data, due to the requirement of having more than one species on either side of the split. We therefore also analyzed the fate of duplicated genes. In this case, we tested whether duplication products are more similar if the non-duplicated members of the gene family have low expression distances across the species outside the duplication event. Indeed, we found significant negative correlations between the median expression distance among the non-duplicated genes and the intra-species correlation of the duplicated genes (Fig. 7B). For example, duplicated genes in fish were more similar (i.e. had a higher correlation) when the corresponding tetrapod genes had more similar expression patterns (i.e. had a low expression distance): r_s_=−0.11 for 2045 pairs of duplicated genes, corresponding to a p-value of 3e-7. Taken together, these two observations implied that for a significant fraction of genes, the rates of change in gene expression patterns were correlated between independently evolving clades.

**Figure 7:**
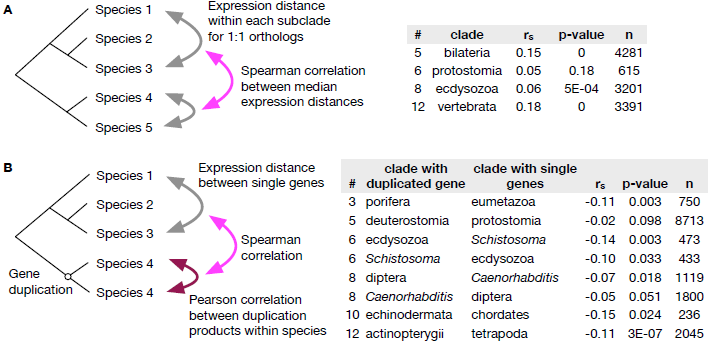
Correlations between expression conservation rates. **A** For 1:1 orthologs, the expression distance across internal nodes was compared. In the dataset, there were only six splits with at least two species on both sides of the split. For example, when genes were similar within tetrapods, they also tended to be similar within fishes. **B** The rate at which gene duplication products diverge was negatively correlated with the expression distance among single-copy genes in related species. Only correlations with p-values below 0.1 are shown in this table. (# – Number of clade in Fig. 5, n – count of 1:1 orthologs [A] or duplicated genes [B])

## Evolution of the Beta Catenin Protein Family

We seleteced the beta catenin protein family [48] as an example to illustrate the implications of our work. Beta catenin proteins are involved in regulating cell adhesion and gene transcription through the Wnt signaling pathway. Ancestrally, there was a single beta catenin protein, which duplicated independently in the nematode and vertebrate lineages [49]. Hence, *Drosophila, Anopheles* and *Schistosoma* only have one beta catenin, armadillo. We found this protein to be similar in its expression patterns with both the vertebrate and nematode beta catenins (Fig. 8), which is indicative of their functional similarities [50]. In vertebrates, two forms exist: beta catenin and plakoglobin. These two proteins have largely overlapping functions [51] and consequently, their observed expression distance was very low. In nematodes, the outcome of the repeated gene duplications [52, 53, 54] is very different: three of the duplication products (*hmp*–2, *wrm*–1, and *sys*–1) are very similar to each other in their expression patterns, which can be explained by their cooperation in in the non-canonical Wnt signaling pathway and the SYS pathway [55]. These three proteins had high expression distances to *bar*–1. In contrast to them, *bar*–1 is part of a canonical Wnt signaling pathway [55]. We also observed that *bar*-1 had a low expression distance to the vertebrate plakoglobin, while *hmp*–2, *wrm*–1, and *sys*–1 had high expression distances. Among the nematode genes, vertebrate beta catenin had the lowest expression distance with *hmp*–2. This example illustrates that our method is able to uncover patterns of functional similarity and divergence both between closely related species and across large evolutionary distances.

**Figure 8:**
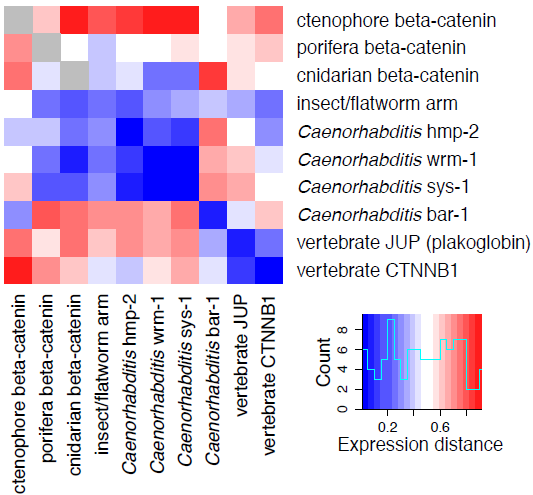
Expression similarity and divergence in the beta catenins. For each group of genes, the median expression distance is shown (see (Fig. S10 for individual expression distances). The unduplicated beta catenins from insects, flatworms and cnidarians a similar to all other protein groups, while functional and expression divergence has occurred independently among nematodes and vertebrates.

## Discussion

The presented analysis established and benchmarked a new method, and provided two examples of biological conclusions that can be reached with our method: there is widespread conservation of expression regulation across very large evolutionary distances, and the expression programs of different gene families evolve at distinct rates. Presumably, the latter observation is explained by variable functional constraints between gene families.

In particular, we have shown that tissue-specific gene expression can be predicted across large evolutionary distances, even in the absence of apparent similarities between the species’ tissues. Our approach can be rationalized as follows: we assume that evolution conserves the coexpression of functionally related genes, both on the level of homologous cell types and on the level of functional modules that occur in unrelated tissues. Our analysis demonstrated that the expression patterns of such conserved gene modules can be predicted across species using 1:1 orthologs as “anchors.” This approach worked despite the fact that the tissues themselves are only conserved within smaller clades. Control of gene expression by transcription factors, miRNAs and other factors is known to turn over rather quickly [56, 57, 58]. Most probably, functional dependencies between genes lead to shared expression patterns over large evolutionary distances. Further research will be needed to reveal which expression similarities between tissues are caused by homology and which are caused by convergent evolution.

## Methods

### Detection of Orthologous Proteins

To determine orthology relations between genes, we assembled groups of orthologs (OGs) using the eggNOG pipeline [59] on the genomes of the choanoflagellate *Salpin-goeca rosetta* and 67 animals. We then computed gene trees for all OGs using GIGA [60], which we then analyzed to extract 1:1 orthologs and duplication events.

### Expression Data Pre-processing

Datasets were obtained either from repositories like ArrayExpress and GEO, from supplementary materials or the respective websites of the resources. Expression profiles were then mapped to our set of genes by one of the following methods (see Table S1): If possible, genes were mapped by given identifiers, such as Affymetrix, Ensembl or WormBase identifiers. If identifiers could not be used for microarrays, we mapped probe sequences to transcripts using exonerate [61], allowing for up to three mismatches and discarding probes that mapped to multiple genes. In the case of RNA-seq data without matching identifiers, we trimmed adapters and mapped reads to annotated transcripts using tophat2 and cufflinks 2.1.1 [62, 63] and used the resulting FPKM counts.

In initial small-scale tests, we tested several normalization methods [11, 64], and settled on a z-like normalization of expression vectors **x**, which corresponds to the Euclidean normalization of **x** minus its median value 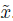.

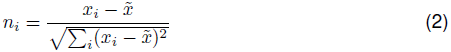

RNA-seq data, e.g. the *Drosophila* modENCODE dataset, contained zeros, which were of course not suitable for logarithmic analysis. For these datasets, we determined the expression value of the 1/1000^th^ quantile of all genes with non-zero expression. All expression values were incremented by this value.

### Mapping of Tissue Expression Patterns

For each pair of datasets, individual linear models were fitted for each tissue of the target species, using the tissues of the source species as input. (Note that due to the normalization, one tissue is redundant and therefore left out. This also implies that the coefficients of the linear model are not directly interpretable.) The set of 1:1 orthologs between the two species was used as to fit the linear models. When there were multiple probes per gene, all combinations of probes were added to the tissue expression matrix. When there were many tissues in the source species, but few 1:1 orthologs, there was the danger of over-fitting. We therefore allowed only one predictor (i.e. one tissue from the source species) per 15 samples (i.e. 1:1 orthologs) [65]. For each pair of species, the safe number of predictors was calculated. If there were too many tissues, we combined tissues using k-means clustering and used the centers of the clusters as predictors. This situation only occurred for six out of 1260 dataset pairs. The fitted models were then applied to all genes of the source species, yielding corresponding predicted expression patterns in the target species. Since 1:1 orthologs are used for training, we used predictions from a 10-fold cross-validation for these genes.

### Mathematical Description of Expression Mapping

To illustrate our approach, we describe it for a specific pair of datasets, namely mapping expression values from fly to mouse (dataset “MOE”). The same procedure can be applied to all pairs of species. To predict tissue expression patterns of the 51 mouse tissues based on the 26 fly tissues, we fitted 51 separate linear models for each mouse tissue based on 1:1 orthologs.

A given dataset of gene expression values across many tissues of a species can be treated as an expression matrix: Rows correspond to genes and tissues to columns. Hence, it is possible to look at *gene expression vectors* that correspond to a single row, and *tissue expression vectors* that correspond to a single column. Consider the matrices of normalized expression values for fly F^0^ and mouse M^0^. F^0^ contained 13,264 rows corresponding to 12,225 genes and 26 columns. M^0^ contained 23,624 rows for 14,307 genes and 51 columns. From the 3120 1:1 orthologs, sub-matrices F and M were constructed such that the same row in the two matrices corresponds to a given pair of 1:1 orthologs. When multiple expression measurements per gene were available, the matrices contained all possible combinations of measurements. (E.g. if there were three probes corresponding to one gene, and two for the ortholog, a total of six rows were dedicated to this pair of orthologs.) Due to these combinations, F and M each had 4447 rows. A single linear model to fit expression values in mouse tissue *t* for genes *g* was thus found by minimizing the errors *∈*:

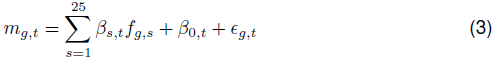

Only 25 parameters were needed in the sum, because the normalization produced a matrix with equal row sums. Therefore one variable was redundant. This approach can also be formulated as a matrix multiplication, using **B** = [*β_s,t_*] as parameter matrix, **B**_0_ = [*β*_0_,_*t*_] for the offsets, and **E** = [ε*_g,t_*] as error matrix:

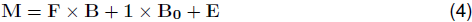

Once **B** and **B**_0_ have been determined, they can be applied to the full expression matrix **F**^0^ to create a matrix **V** of virtual expression values for fly genes in mouse tissues:

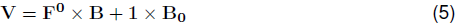

### Cross-species Correlations Between Expression Patterns

For each fly gene *x* with its corresponding gene expression vector 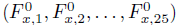 an expression vector based on mouse tissues had been predicted: *X* = (V*_x_* _1_,V*_x_*, …, V*_x_*,_51_).

Thus, for any mouse gene *y* with expression vector 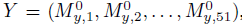, the weighted sample Pearson correlation coefficient *r_x y_* could be calculated (Fig. 2A):

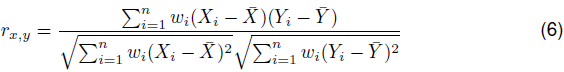

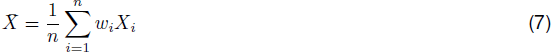

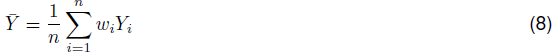

Weights on the tissues were calculated using the Gerstein-Sonnhammer-Chothia (GSC) weighting scheme to reduce the effect of uneven coverage of different anatomical regions [66]. For example, in the mouse tissue dataset, there were many different brain tissues with highly correlated expression patterns. Hence, a gene that was well predicted in one brain tissue was likey to be well-predicted in other brain tissues. When multiple measurements were available for the source or target gene, we reported the maximum of all pairwise correlations.

### Computation of Expression Distances

For each pair of datasets, we computed a matrix of predicted expression patterns of all genes from the source species. We observed a strong correlation between the cross-species expression correlation and the number of coexpressed genes in the target species (Fig. S3). This strong correlation indicated that predictions were biased towards the average target gene (i.e. the average expression profile of all genes considered in the target species), which in turn was similar to many target genes. As a consequence, these “close-to-average” target genes had higher correlations with mapped source genes, and thus seemed more conserved. To counter this effect, target genes were split into ten bins according to the number of coexpressed genes in the target species (Fig. 9). For each bin, we separately determined the distribution of cross-species expression correlations between all genes. Given this distribution, we determined a conversion function from the cross-species expression correlation to the corresponding quantile.

Thus, there exist ten conversion functions from weighted Pearson correlation to an uncorrected expression distance. For a given pair of genes, the final expression distance is interpolated from the two adjacent bins. We determined the number of coexpressed genes for each target gene as follows: we first computed all pairwise correlations among the target genes of the training set. Then, we determined the correlation cutoff corresponding to the top 10%, and counted for each gene how many other target genes were among the global top 10% correlations. For technical reasons, we sampled one million pairs of background genes, such that the lowest possible expression distance is 1e-6.

**Figure 9:**
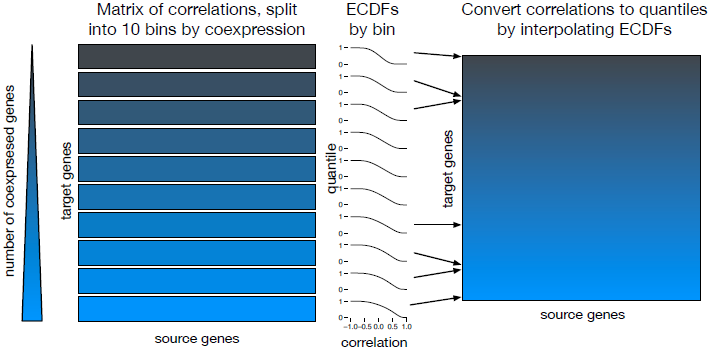
Conversion from correlations to expression distances. For each pair of datasets, all pairwise Pearson correlations between the actual expression patterns and the mapped patterns were computed. This complete matrix of correlations was then split into ten parts, according to the number of coexpressed target genes. For each bin, we separately calculated the ECDF, i.e. the conversion function between correlations and quantile. To retrieve the expression distance of a given pair of source and target genes, the number of coexpressed genes for the target gene was used to select the two closest bins and their ECDFs. To reduce the effect of small differences in the number of coexpressed genes, the expression distance was then computed by interpolating the quantiles returned by the two ECDFs. (Edge cases, where only the first or last bin are appropriate, were treated separately.)

## Data Access

Protein sequences, normalized datasets, assignments and expression distances of 1:1 orthologs have been deposited at http://dx.doi.org/10.6084/m9.figshare.1362211. Separately, all pairwise mappings of expression patterns between datasets are available at http://dx.doi.org/10.6084/m9.figshare.1362240

## Acknowledgements

The authors thank Anthony A. Hyman and Vineeth Surendranath for helpful discussions.

## Author Contributions

AB and MK conceived the study, planned the analyses and wrote the paper. MK conducted all analyses.

## Supplementary Information

**Figure S1:**
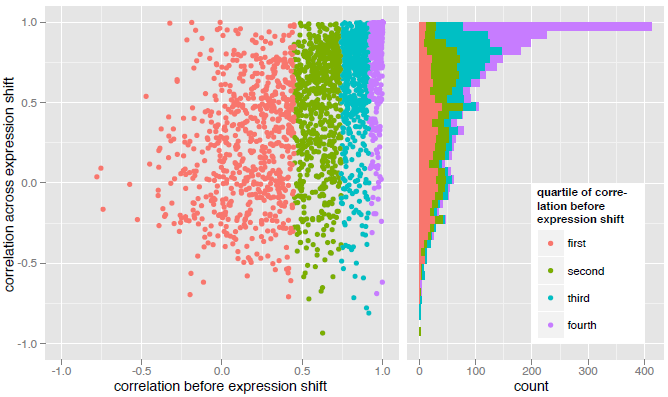
Lineage-specific expression shifts do not change expression patterns. Proteins that have been reported to have lineage-specific changes in expression, e.g. between primates and non-primate species [10] have highly correlated expression patterns even across the expression shift if the expression pattern has been fixed before (fourth quantile, purple).

**Lineage-specific Expression Shifts and Relative Expression Patterns**

In the main text, we investigated changing and conserved expression patterns. A previous analysis of expression patterns in six tissues across eight mammals and chicken concluded that while the expression of most genes is under purifying selection, there are also many cases of lineage-specific expression shifts [10]. However, in a re-analysis of this data, we found that these changes occurred mainly on an absolute expression level and that even across the expression shifts, the expression patterns which were reported in the original data set stayed highly correlated (Fig. S1): For the set of genes with significant expression shifts, we found a median correlation of 0.68 between the expression patterns of the species with the expression shift and the species with unchanged expression (“outgroup”). We suspected that for some genes, the expression pattern only becomes fixed after the expression shift. Indeed, when we divided the genes into quartiles according to the median correlation within the set of proteins in outgroup, we found that in the bottom quartile the median correlation across the expression shift is 0.25, while in the top quartile the median correlation is 0.95. In other words, once an expression pattern becomes fixed, it is retained even across lineage-specific expression shifts.

**Figure S2:**
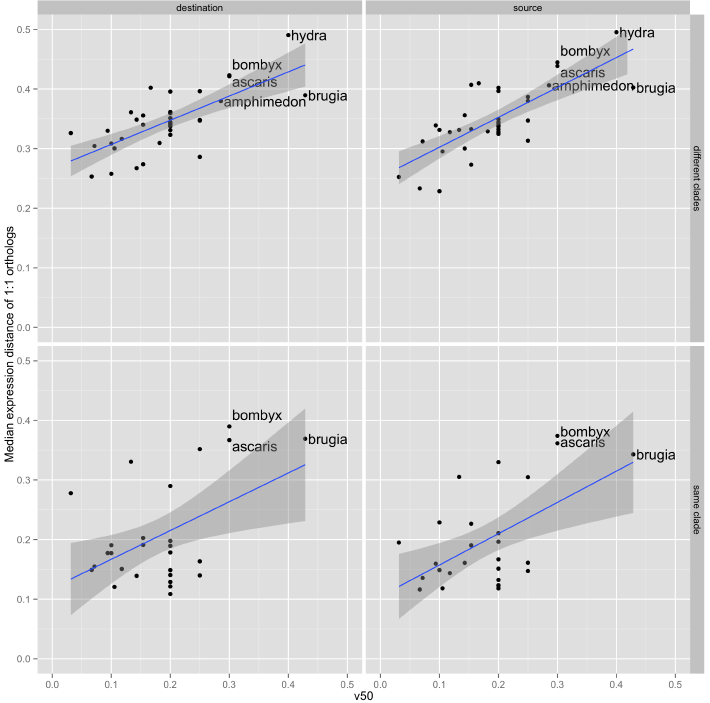
Correlation between dataset quality and mapping quality. For each pair of datasets, the median expression distance of 1:1 orthologs (Fig. 4) can be treated as the mapping quality. For each dataset, we then calculated the median mapping quality over all dataset pairs for which the given dataset is either the source or target dataset. We further distinguish between dataset pairs of the same clade (e.g. two vertebrates) versus pairs from different clades. (Dataset pairs within a clade are only considered for clades of at least three species.) In all combinations, there is a correlation between V50 and the mapping quality. We therefore use this measure to exclude five datasets. (Blue line: linear fit; shaded area: 95% confidence interval.)

**Properties of Proteins that Influence the Mapping**

It is to be expected that properties of the considered genes have an effect on how well the genes’ expression patterns can be mapped. For instance, it seems likely that genes that are well-conserved on the protein sequence level should also have conserved expression patterns. Conversely, 1:1 orthologs may appear to have dissimilar expression patterns either due to biological reasons (e.g. functional divergence) or due to technical reasons (e.g. measurement noise, inability to map the expression pattern correctly). We therefore tested eight different properties to which extent they are correlated with expression similarity (using Spearman’s rank correlation coefficient). The tested properties were: number of (same-species) proteins with similar expression pattern, degree in the STRING 9.1 protein-protein interaction network (using experimental and text-mining evidence and a confidence score threshold of 0.5) [67], number of isoforms (according to data from Ensembl, WormBase and FlyBase), number of residues, tissue specificity [68], absolute expression level, sequence similarity between the considered proteins, and pleiotropy (for mouse proteins [69]). Almost all properties had a significant influence (Fig. S3). The effects of sequence similarity, total expression level and degree were consistent with previous findings that these factors are inversely correlated with gene loss [70].

**Figure S3:**
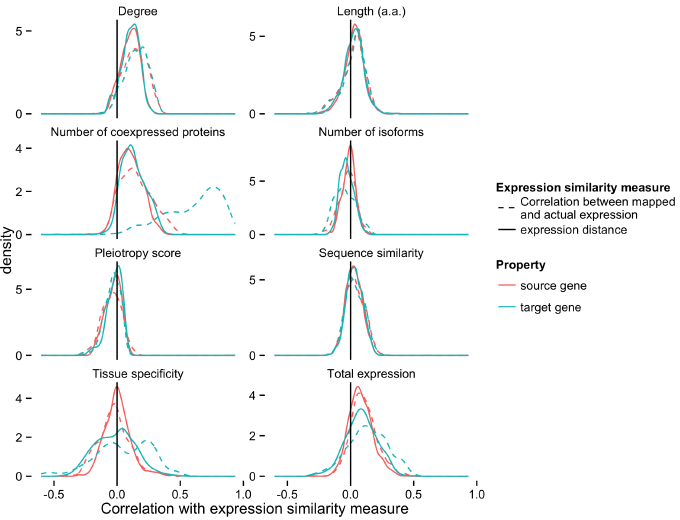
Gene properties correlated with expression similarity. Different properties of the source (red) or target gene (blue) influenced the distribution of expression distances. To measure this influence, we computed the correlation between the gene properties and the expression similarity of 1:1 orthologs. When the correlation between mapped and actual expression patterns was used as the expression similarity (dashed lines), there was a very high correlation with the number of coexpressed target genes. That is, when a target gene had many genes with similar expression patterns, then the expression correlation with its 1:1 ortholog tended to be high. Correcting for this (solid lines), this correlation became lower. Genes corresponding to proteins with high degree (i.e. number of interactions) could be mapped better, while target genes with many isoforms resulted in a worse mapping.

**Figure S4:**
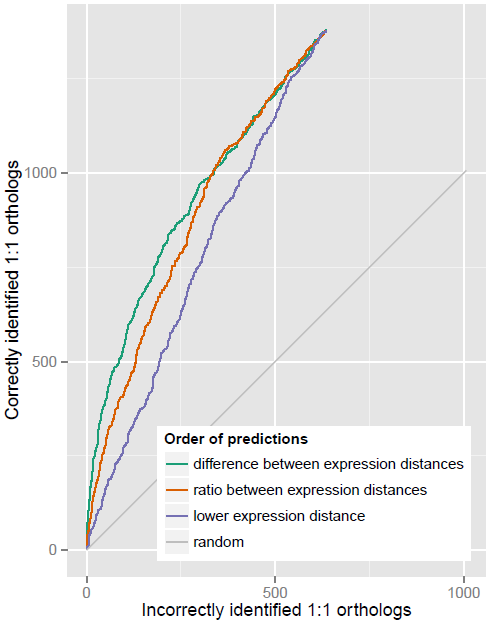
Prediction of 1:1 orthologs from best hits. For each 1:1 ortholog between fly and *C. elegans,* the source genes expression pattern was mapped to *C. elegans* and compared to the top two BLAST hits. If the mapped expression pattern was more similar to the actual ortholog, it was counted as correctly identified. Thus, a perfect prediction method would be a vertical line. Ordering the predictions by the difference between the two expression distances was the most successful strategy.

**Benchmark 1: Identification of 1:1 Orthologs**

In a first benchmark, we tested whether the expression similarity could be used to identify 1:1 orthologs from top BLAST hits. For each dataset pair, we used BLAST to find the top two hits for each protein of the source species, discarding proteins with only one hit. After training the expression mapping on an independent set of genes as outlined above, we then computed the expression similarities for the top two hits, and checked whether the gene with the lower expression distance corresponded to the actual 1:1 ortholog. For example, mapping from fly to *C. elegans,* 67.5% of 2014 one-to-one orthologs could be correctly identified (p-value of Binomial test: 5e-57; median across all dataset pairs: 60%). Predictions could be ordered in different ways according to the expression distances between the two pairs of genes: by the lowest expression distance, by the difference of the expression distances or their ratio. Of these, the difference between the expression distances performed best in distinguishing confident predictions from less confident predictions (Fig. S4).

**Figure S5:**
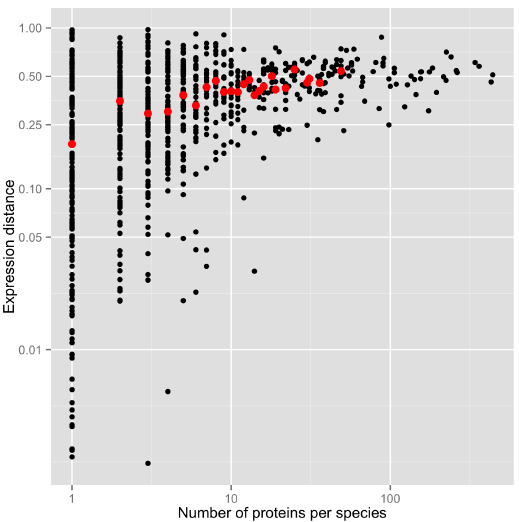
Correlation between expression distance and shared protein folds. Proteins that belong to structural families with few members are more similar in their expression patterns than proteins from large families. Red dots denote the median when at least five superfamilies have the same number of proteins per species. Here, the mapping from fly to *C. elegans* is shown.

**Benchmark 2: Analysis of 3D Protein Structure**

As a further test, we checked if genes corresponding to proteins with the same structure were more likely to have lower expression distances than unrelated proteins. Using the Gene3D database [45], we determined CATH folds for all proteins that we could map to the database (resulting in 15 species and 23 datasets). For each dataset pair, we then analyzed each homologous superfamily, computing the median expression distance for all proteins of the superfamily. The superfamilies contain varying numbers of proteins, and we found a correlation between the expression distance and the size of the superfamilies (Fig. S5): Those with many members (and thus more different functions) had more diverse expression patterns. For example, mapping fly to *C. elegans,* the Spearman correlation between the number of proteins per species (using the maximum of the two species) and the median expression distance was 0.40. Between human and mouse (GNF dataset), the Spearman correlation was 0.46. Across all dataset pairs, the median Spearman correlation was 0.28.

**Benchmark 3: Phenologs**

Finally, we used functional information to evaluate our method. We applied the phenolog concept [46] to validate that genes from different species with similar tissue expression are functionally related. Based on orthologous genes, related pairs of functional annotations (Gene Ontology terms, FlyBase and WormBase phenotypes) are predicted by looking for significant overlap between OGs that correspond to the functional annotations. This leads to phenologs, i.e. pairs of functional annotations with a certain p-value that represents their cross-species similarity. For each pair of well-annotated species (mouse, human, fly, *C. elegans),* we tested all OGs excluding 1:1 orthologs. For each OG, we found the phenolog with the lowest p-value. For all cross-species gene pairs in this OG, we then determined their expression distance and whether their functional annotation matched the phenolog pair.

**Figure S6:**
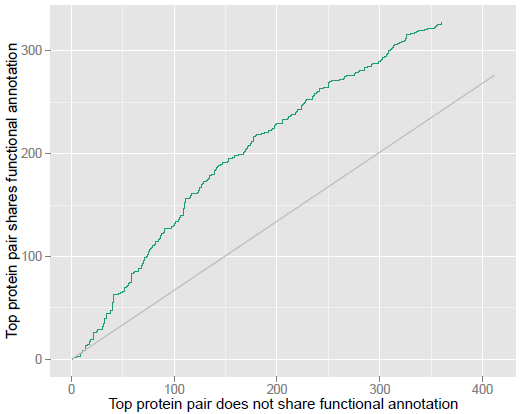
Benchmarking based on phenologs. For each OG between fly and *C. elegans,* we selected the phenolog with the lowest p-value. We then tested whether the gene pair with the lowest expression distance shared the functional annotation predicted by the phenolog. Predictions were ordered by the difference between the lowest and second lowest expression distance. Randomly choosing gene pairs from the OGs results in the grey line.

First, we noted that the distributions of expression distances differed between gene pairs with matched and mismatched annotations: For fly and *C. elegans,* the Wilcoxon rank sum test p-value was 6e-19 (median p-value across all dataset pairs: 2e-9). Second, for each OG, we looked at the gene pair with the lowest expression distance and checked if both genes matched the expected functional annotation based on the phenolog. We ordered OGs by the difference between the lowest and second lowest expression distances. Mapping fly to *C. elegans,* 47.7% of all top predictions had matching functional annotations, compared to an expected fraction of 40.2% (Fig. S6). This corresponds to a relative increase of 19% over the expected fraction. Between human and mouse (GNF dataset), this increase is 34%. The median increase among all dataset pairs is 10%.

**Figure S7:**
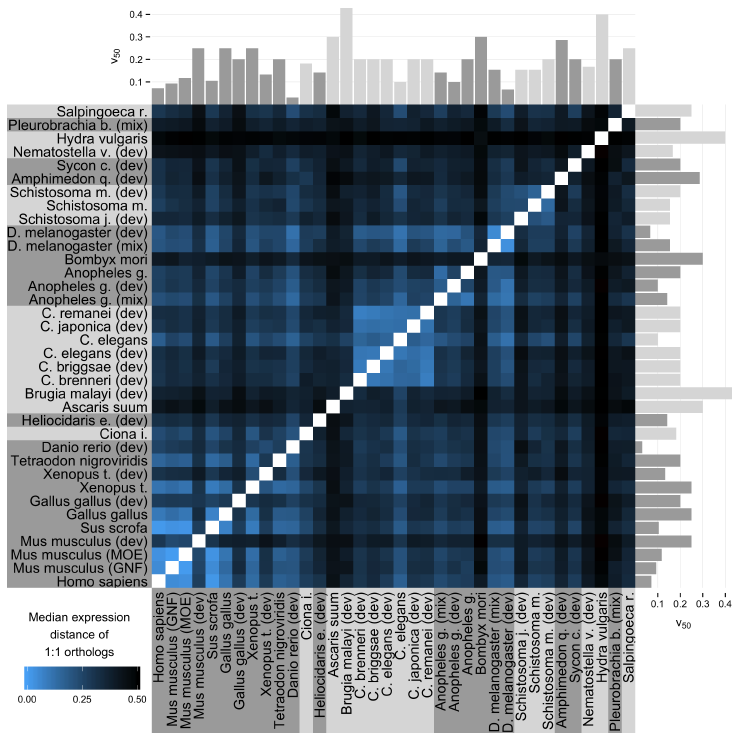
Conservation of expression patterns throughout the metazoans and choanoflagellates. In this version of Fig. 4, all datasets are shown without any filters for dataset quality. Therefore, five additional datasets are shown: *Hydra vulgaris, Amphimedon queenslandica, Bombyx mori, Brugia malayi* and *Ascaris suum.* For these species, the quality of the datasets prevented better mapping performance.

**Figure S8:**
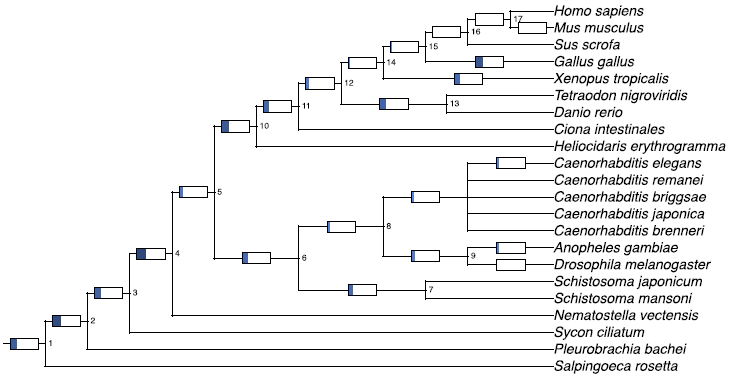
Most conserved expression across animal clades. As in Fig. 5, the median expression distances of 1:1 orthologs are shown. However, instead of the median across all datasets, the median expression distance of the best dataset pair is used. Charts at species branches show how well expression patterns could be mapped between different datasets of the same species.

**Figure S9:**
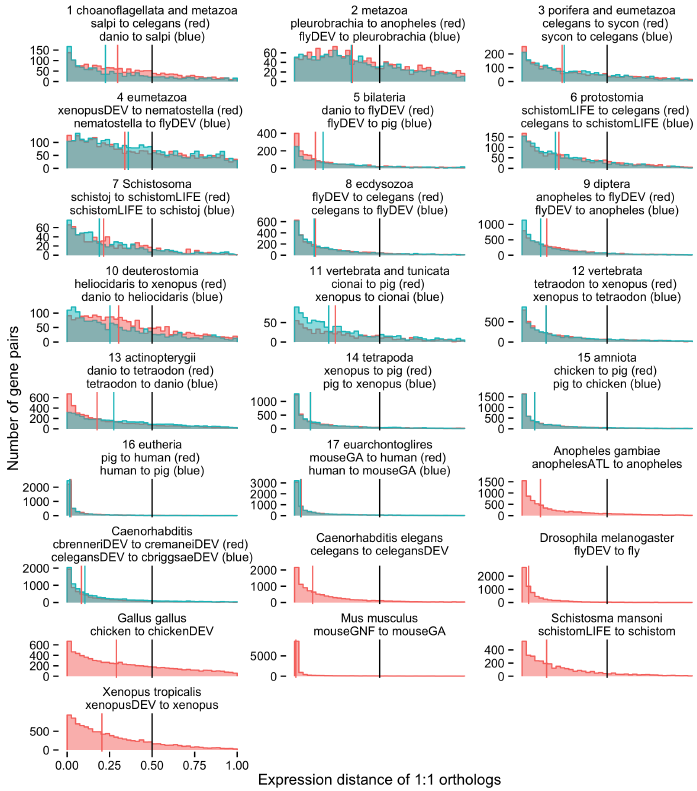
Distribution of conserved expression for best dataset pairs. For each internal node, the distribution of expression distances is shown for the datasets given. These datasets show the highest degree of conservation for the respective internal node. See Table S1 for a description of the abbreviated dataset names.

**Figure S10:**
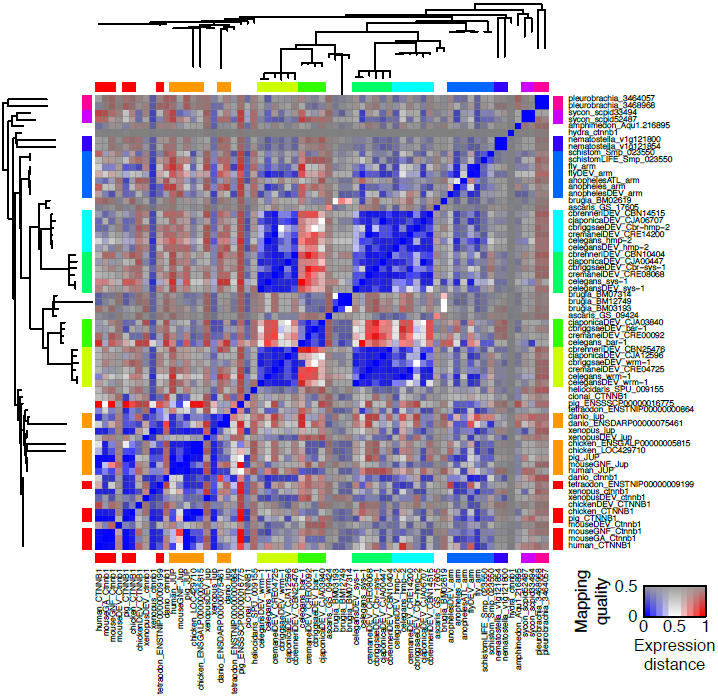
Expression distances of beta catenins. The heatmap shows the expression distances of beta catenins throughout the animal kingdom as color scale. Individual cells are shaded in grey according to a measure of mapping quality of the pair of datasets, namely the median expression distance of 1:1 orthologs (Fig. 4). Genes were clustered according to the gene tree built from a multiple sequence alignment. The colored band along the edge of the heatmap denote the ten groups of proteins which have been analyzed in Fig. 8. Genes that belonged to low-quality datasets or showed divergent expression patterns to their closest neighbors were discarded.

